# nPIST: A Novel Actin Binding Protein of trans-Golgi Network

**DOI:** 10.1101/270363

**Authors:** Swagata Das, Priyanka Dutta, Mohit Mazumder, Soma Seal, Kheerthana Duraivelan, Dibyendu Samanta, Samudrala Gourinath, Sankar Maiti

**Affiliations:** Department of Biological Sciences, Indian Institute of Science Education and Research, Kolkata. Nadia, West Bengal, INDIA; School of Life Sciences, Jawaharlal Nehru University, New Delhi, INDIA; School of Bioscience, Indian Institute of Technology Kharagpur, Kharagpur, West Bengal, INDIA

**Keywords:** nPIST, trans-Golgi network, WH2-domain, actin, vesicular trafficking

## Abstract

npist is the neuronal isoform of PIST, a trans-golgi associated protein involved in major modulation of vesicular trafficking. nPIST interacts with glutamate delta2 receptor (GluRδ2) in Purkinje cells. Our study shows nPIST as a novel actin binding protein. Our structure based sequence analysis shows nPIST contains one WH2-like domain. Further our experimental analysis illustrates that fragment of nPIST consisting of WH2-like domain binds to actin. Moreover it was found that nPIST contains several regions involved in interaction with actin. The binding of nPIST to actin through multiple actin binding regions facilitated actin filament stabilization *in vitro*. *In vivo*, nPIST localized actin in perinuclear region as a blotch when ectopically expressed.

## Introduction

PIST (PDZ domain protein interacting specifically with TC10) also known as GOPC (Golgi-associated PDZ and coiled-coil motif-containing protein) is primarily discovered as TC10 Rho GTPase interacting protein (Neudauer *et al.*; 2001). It is also known as CAL [cystic fibrosis transmembrane conductance regulator (CFTR)-associated ligand] (Cheng *et al.*; 2002, and 2004) and FIG (Fused in Glioblastoma) (Charest *et al.*; 2003, and Cheng *et al.*; 2004). PIST is expressed in all type of mammalian tissues whereas its neuronal isoform known as nPIST is exclusively expressed in mammalian whole brain preferentially in parallel fiber synapses of cerebellar Purkinje cells, hippocampus and cerebral cortex (Cuadra *et al.*; 2004, Yue *et al.*; 2002, Yao *et al.*; 2001, and Chen *et al.*; 2012). It has two N-terminal coiled-coil domains and one C-terminal postsynaptic density-95/discslarge/zonula occludens-1 (PDZ) domain (Neudauer *et al.*; 2001, Cuadra *et al.*; 2004, Yue *et al.*; 2002, and Yao *et al.*; 2001). The second coiled-coil domain is required for its golgi localization and interaction with Golgi resident proteins (Charest *et al.*; 2001, and Hicks and Machamer; 2005) whereas the PDZ domain is known to interact with several plasma membrane destined proteins. In the past decade, PIST has been well established as a key Golgi resident candidate for regulation of intracellular sorting and trafficking of plasma membrane targeted proteins.

In initial studies, PIST emerged as interacting partner of several transmembrane proteins having wide array of functional aspect. Interestingly, the mode of action of PIST for different proteins differs from one another. The plasma membrane expression of Rhotekin, mGluR5a (Metabotropic Glutamate Receptor subtype 5a), and Frizzled5 is enhanced by their interaction with PIST (Ito *et al.*; 2006, Cheng *et al.*; 2010, and Yao *et al.*; 2001). On the other hand, PIST restricts plasma trafficking of diverse types of membrane spanning proteins. Among all these proteins, some are ion channels like CFTR, CIC3B (ClC Chloride Channel 3B) (Cheng *et al.*; 2002, 2004, and Gentzsch *et al.*; 2003); some are receptors like SSRT5 (Somatostatin Receptor subtype 5), mGluR1a (Metabotropic Glutamate Receptor subtype 1a), ß1AR (ß1-Adrenergic Receptor), ß2AR (ß2-Adrenergic Receptor), CRFR1 (Corticotropin-releasing Factor Receptor 1) (Wente *et al.*; 2005, Zhang *et al.*; 2008, He *et al.*; 2004, Yang *et al.*; 2015, and Hammad et.; 2015); cell adhesion molecules cadherin23 (Xu *et al.*; 2010), and protocadherin15 (Nie *et al.*; 2016); tight junction proteins claudin1 and 2 (Lu *et a1.*; 2015); ABC transporter mrp2 (Li *et al.*; 2017) etc. The Golgi retention of most of these proteins results in their degradation (Cheng *et al.*; 2010, 2013, and Lu *et al.*; 2015) but in contrast, ß1AR is stabilized and mGluR5a is salvaged from polyubiquitination through interaction with PIST (He *et al.*; 2004, and Cheng *et al.*; 2010).

Remarkably, PIST is not only involved in golgi mediated trafficking but might also play major role in golgi morphology and maintenance. In PIST knockdown mice, spermatogenesis gets completely hampered and develops several cellular process dysfunctions which cumulatively results in globozoospermia. Among these cellular deteriorations, the major event is malformation of acrosomal cap in spermatids. Other than occurrence of vesicular or fragmented acrosome; nuclear malfunction and abnormal arrangement of mitochondria also takes place. Moreover, in PIST^−/−^ condition seminiferous tubule loses its normal F-actin filament distribution. Actin filaments fail to surround the adjacent spermatids and get accumulated in basal region (Yao *et al.*; 2002, and Ito *et al.*; 2004). The neuronal isoform nPIST contains eight amino acids extra (151-158 aa) in the second coiled-coil domain. nPIST is primarily known to interact with Glutamate δ2 Receptor (GluRδ2) (Yue *et al.*; 2002). Like its non-neuronal isoform, nPIST is also involved in vesicular trafficking regulation. The clustering of AMPA receptor GluR1 to synapse is synchronized by interaction between AMPAR-interacting protein Stargazin and nPIST (Cuadra *et al.*; 2004).

The indispensable role of both PIST and nPIST in diverse cellular and physiological activities implies that they might have direct or indirect interaction with actin cytoskeleton. Functional characterization of nPIST is not studied in details like that of its non-neuronal isoform. In this study we have characterized nPIST as a novel actin binding protein. As nPIST have mostly coiled-coil structure, we searched for WASP homology 2 domain (WH2) in nPIST by multiple sequence alignment with WH2 domains of other well characterized actin binding proteins. The alignment detected one WH2-like domain having conserved amino acids which are characteristic feature of WH2 domain of several actin binding proteins (Paunola *et al.*; 2002). Our study suggested that nPIST is able to interact with actin through its multiple actin binding regions and stabilizes filamentous actin in *in vitro* condition. Further it was observed that nPIST gets localized to perinuclear region inside HEK293 cells through its N-teriminal coiled-coil region and causes abnormal accumulation of cellular F-actin co-localizing with it.

## Results

### nPIST, a trans-Golgi associated protein have putative WH2 domain

We build several structural models of nPIST protein using Swiss model and modeller. The models shown in these studies were energy minimized after modelling to get a lower energy and more stable conformation. The full-length nPIST is composed of 463 amino acid residues with two domains; N-terminal coiled-coil region and the C-terminal contain PDZ signaling domain. Based on the sequence information we built various models of N and C terminal of nPIST from closely related structures based on the template search with sequence identity of more than 30 percent sequence identity. The top scoring templates with known domains are shown in (Fig. 1A) along with the sequence coverage at the top. The model of coiled coil domain region (34^th^ – 194^th^ aa, and 74^th^ – 204^th^ aa) is shown in (Fig. 1B) and the PDZ domain region models with identified WH2 domain (213^th^ -374^th^ aa, 201^st^ -370^th^ aa, and 215^th^ -368^th^ aa) is shown in (Fig. 1C). The structural model revealed that N-terminal coiled-coil region as predicted (secondary structure) was highly helical compared to extreme C-terminal PDZ domain region which was comparatively disordered and composed of many loop regions. The central helix formed a hinge giving flexibility to the molecule to move. The full length structure based sequence analysis revealed that nPIST contains WH2 domain within the coiled-coil region. The WH2 domain was identified on the basis of sequence conservation, evolutionary function and structure of the domain. Based on structural model we performed sequence based structural alignment of nPIST with all the available crystal structures of WH2 domain. The alignment clearly suggested that coiled-coil domain is present in N-terminus of nPIST and the probable WH2 domain is present around 230^th^ to 250^th^ amino acids (Fig. 1D). nPIST WH2 domain folded exactly same as seen in various structures co-crystallized with G-actin (Dominguez and Holmes, 2011), suggesting that the identified domain might work similarly. To understand more about the structural details of actin binding we modeled nPIST WH2 domain with G-actin using rosetta Flexpepdock server. The docking results suggested that nPIST WH2 domain binds adequately with G-actin (Fig. 1E). The mode of binding is similar to VopL actin complex (Rebowski *et al.*; 2010). The complex was then subjected to further analysis for identification of key residues responsible for actin binding using KFC-2 (Knowledge-based FADE and Contacts) server. The server analyzed several chemical and physical features surrounding an interface residue and predicted the classification of the residue using a model trained on prior experimental data. The final complex obtained after simulations was used as the input for KFC2. The analysis was done for WH2 domain in which TRP-238, GLN-240, LEU-241, ILE-245 were predicted as binding hotspots (Table 1). Figure1F showed location of hotspots at the binding interface of G-actin predicted by KFC2 server.

**Figure 1:**
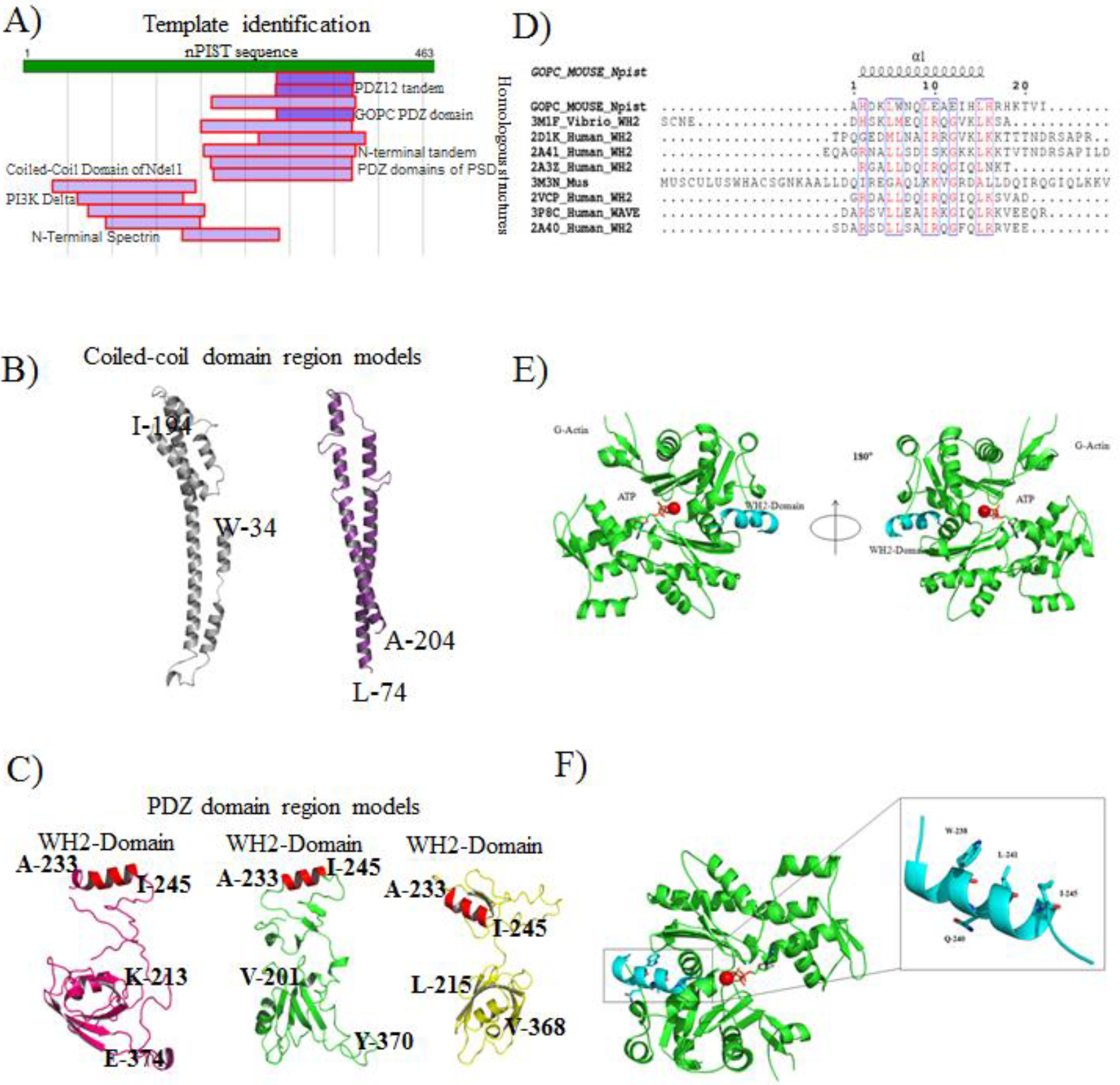
Structure based modelling of nPIST indicated the presence of putative WH2 domain in nPIST. (A) The template identification carried out by Swiss model is shown in figure. The scematics shows the sequence coverage of closely related models available in PDB database with characterized domains labeled B) The models of N-terminal coiled-coil domain region of nPIST C) Models of C-terminal PDZ domain region nPIST with identified WH2 domain (D) The Structure based sequence alignment of WH2 domain sequences obtained from crystal structure of various actin-WH2 domain complexes with nPIST. Binding of probable WH2 domain of nPIST with G-Actin; (E) Two 180° view of the model of G-actin-WH2 Domain complex. (F) The location of hotspots at the binding interface of G-actin predicted by KFC2 server.

**Table 1:**

| Chain ID | Residue | Number | KFC2-A | KFC2-B |
| --- | --- | --- | --- | --- |
| B | ALA | 233 | ----- | ----- |
| B | HIS | 234 | ----- | ----- |
| B | ASP | 235 | ----- | ----- |
| B | LEU | 237 | ----- | ----- |
| B | **TRP** | 238 | Hotspot | Hotspot |
| B | **GLN** | 240 | Hotspot | ----- |
| B | **LEU** | 241 | Hotspot | Hotspot |
| B | GLU | 242 | ----- | ----- |
| B | GLU | 244 | ----- | ----- |
| B | **ILE** | 245 | Hotspot | Hotspot |
| B | HIS | 246 | ----- | ----- |
| B | ARG | 249 | ----- | ----- |

### Full length nPIST has actin binding ability

The full length nPIST (1^st^ - 463^rd^ aa) (Fig. 2A) was cloned in pET28a vector and expressed in *E. coli* Bl21-DE3-RP strain. nPIST was purified as N-terminal 6-His tagged protein (Fig. 2B). Purified nPIST showed molecular size of ~60 KDa on SDS PAGE; however the theoretical size of nPIST is 54 KDa including 6 His-tag flanking region. To resolve this, we had sequenced our cloned nPIST ORF and found that its sequence is identical with the nPIST sequence from database with ID BAC27058.1 from EMBL-EBI. From computational analysis it was predicted that nPIST contains a putative WH2 domain (Fig. 1C and D). To inquire whether nPIST also acts as other WH2 domain containing actin binding proteins, we performed F-actin co-sedimentation assay with full length nPIST. In the reaction set up, increasing concentrations (0.5 μM, 1 μM, 2 μM, 3 μM, 4 μM, and 5 μM) of nPIST was incubated with 5 μM of F-actin. Only 5 μM nPIST containing reaction was used as negative control. After 310 x 1000 g centrifugation, the supernatant and pellet fractions were run in SDS PAGE. The coomassie stained SDS PAGE showed that the band intensity of full length nPIST in pellet fraction gradually increased when added to the reaction in increasing concentration (Fig. 2C). In protein control; there was very negligible amount of nPIST in pellet fraction which was much less than that of the reaction having 5 μM of nPIST incubated with actin. This result illustrated that full length nPIST got co-sedimented with F-actin in the pellet fraction in a concentration dependent manner. This revealed full length nPIST was able to bind actin like other actin binding proteins (Shimada *et al.*; 2004).

**Figure 2:**
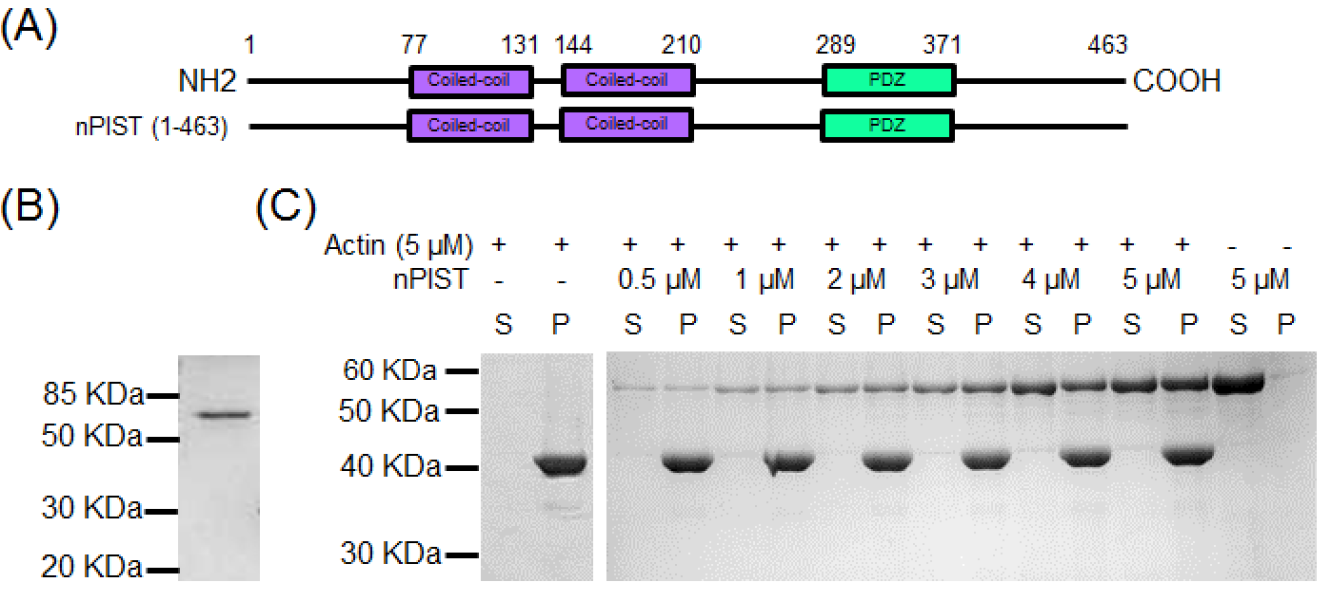
Full length nPIST can bind to actin. (A) Schematic representation of full length nPIST. (B) Purified full length nPIST (1^st^ - 463^rd^ aa) in coomassie stained 12% SDS-PAGE. The protein is expressed in *E. coli* Bl21-DE3-RP cells as N-terminal 6-His tagged and purified using Ni-NTA-agarose bead. (C) SDS-PAGE of supernatant (S) and pellet (P) fraction of actin co-sedimentation assay with 0.5 - 5 μM of full length nPIST.

### nPIST has high affinity for F-actin, but not G-actin

The specific binding of nPIST to actin molecules was determined by Surface Plasmon Resonance (SPR) analysis which monitors molecular interaction in real time condition. Varying concentrations of F-actin and G-actin solutions were passed over purified nPIST protein (immobilized onto a series S CM5 sensor chip) and binding was detected by changes in RUs over time. The association time of both monomeric and filamentous actin to nPIST indicated rapid intermolecular interaction. However, the affinity of F-actin to nPIST was much higher than that of G-actin molecules (Fig. 3, and S1E). In case of F-actin, the equilibrium dissociation constant (*K*_D_) was calculated to be ~ 13.3 nM that illustrates high molecular affinity, with *k*_a_ = 9.25 × 10^4^ M^−1^ s^−1^ and *k*_d_ = 1.23 × 10^−3^ s^−1^ (Table 2). On the other hand, G-actin did not show stable interaction with nPIST (Fig. S1E).

**Figure 3:**
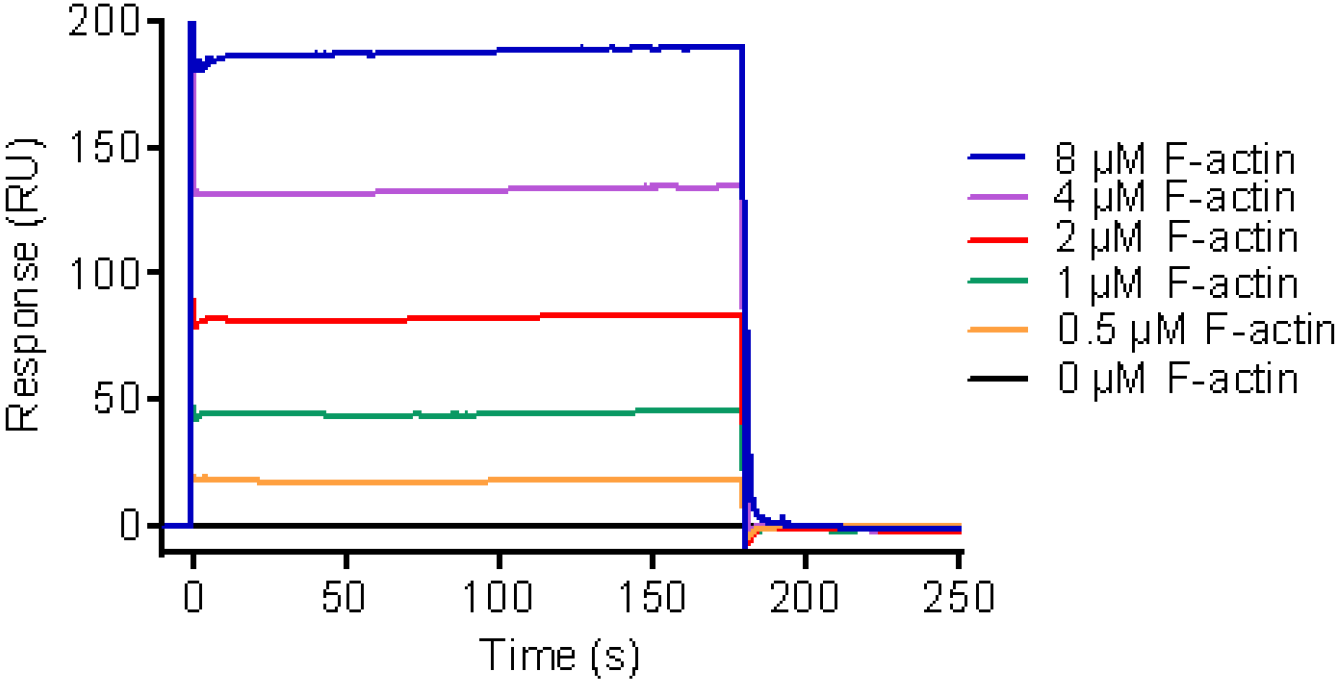
SPR interaction analysis for F-actin binding to nPIST. The sensorgram shows association and dissociation phases of varying concentrations of F-actin (analyte) binding to nPIST (immobilized ligand). The interactions follow rapid kinetics. Fitting the data to 1:1 interaction model gives a *K*_D_ of ~13.3 nM for F-actin.

**Table 2:**

| $K_D$ (M) | $k_a$ (M<sup>-1</sup> s<sup>-1</sup>) | $k_d$ (s<sup>-1</sup>) |
| --- | --- | --- |
| $13.3 \times 10^{-9}$ | $9.25 \times 10^4$ | $1.23 \times 10^{-3}$ |

### WH2 domain containing fragment of nPIST enough for actin interaction

To check whether WH2 domain containing nPIST fragment was solely binding to actin, we had dissected nPIST into two fragments. For that, full length nPIST was sub-cloned in two parts; N-terminal coiled coil region having WH2 domain (CC-WH2 1^st^ - 267^th^ aa) and C-terminal PDZ domain region devoid of WH2 domain (ΔWH2, 268^th^ - 463^rd^ aa,) (Fig. 4A). The CC-WH2 and ΔWH2 fragments were purified as 6-His tagged protein (Fig. 4B). The ΔWH2 fragment was visualized in SDS PAGE at ~30 KDa (Fig. 4B, lane 2) but the theoretical size of the protein is 24.5 KDa with tag. We had sequenced our construct and like full length nPIST, ΔWH2 fragment also showed sequence identity with the database sequence. F-actin co-sedimentation assay with these fragments showed CC-WH2 fragment was co-sedimented along with F-actin in a concentration dependent manner in pellet fraction when 4 μM, 8 μM and 12 μM of CC-WH2 fragment was used for the reactions (Fig. 4C). Surprisingly, the ΔWH2 fragment was also visible in pellet fraction with sedimented actin in increasing amount (Fig. 4D). In both the cases, there were very negligible amount of CC-WH2 and ΔWH2 protein in the pellet fraction of the respective protein control reactions. From this result, it was palpable that CC-WH2 as well as ΔWH2 fragment of nPIST had the ability to bind to actin.

**Figure 4:**
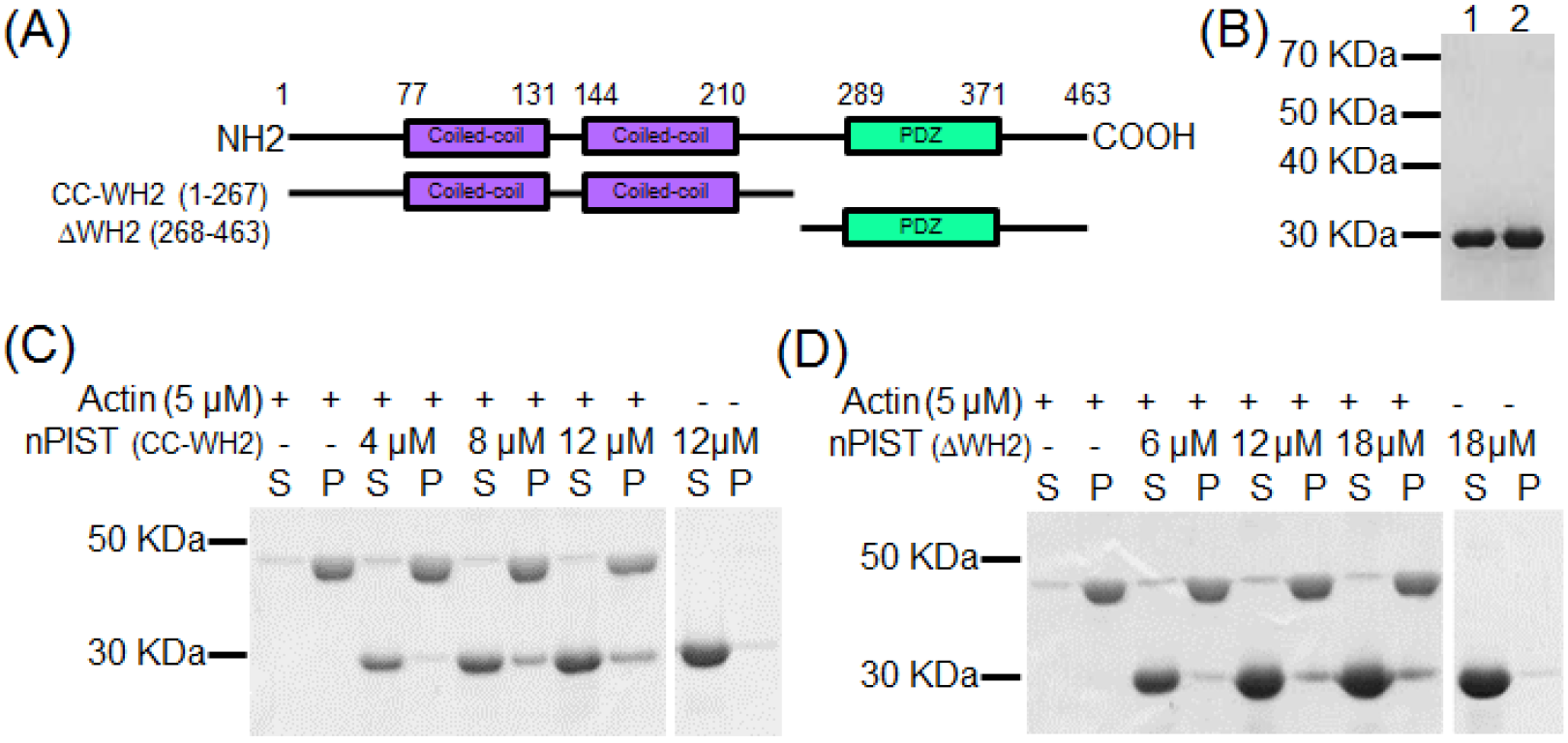
Fragment of nPIST lacking WH2 domain also binds to actin. (A) Illustrative portrayal of fragments nPIST showing N-terminal coiled-coil domain containing construct (CC-WH2) and C-terminal PDZ domain containing construct (ΔWH2). (B) SDS-PAGE of 6-His tagged purified protein of CC-WH2 (lane 1) and ΔWH2 (lane 2) fragments of nPIST. Supernatant (S) and pellet (P) fraction of actin co-sedimentation assay of CC-WH2 (C) and ΔWH2 (D) fragments separated by 12% SDS-PAGE and stained with coomassie.

### nPIST interacts with actin through multiple regions

The above result hinted that nPIST might have more sites engaged in interaction with actin other than the putative WH2 domain (Fig. 1B). To identify other possible actin binding regions, full length nPIST was truncated into smaller stretches and these fragments were sub-cloned in pET28a vector. The fragments containing first coiled-coil domain (CC1, 1^st^ - 138^th^ aa), second coiled-coil domain (CC2, 140^th^ – 210^th^ aa), modeled WH2 domain (WH2, 200^th^ - 289^th^ aa) and only PDZ domain (PDZ, 285^th^ – 463^rd^ aa) were taken into consideration (Fig. 5A). The N-terminal 6-His tagged CC1 fragment (1^st^ - 138^th^ aa) (Fig. 5B), CC2 fragment (140^th^ – 210^th^ aa) (Fig. 5C), WH2 fragment (200^th^ −289^th^ aa) (Fig. 5D), and PDZ fragment (285^th^ - 463^rd^ aa) (Fig. 5E) were overexpressed in *E. coli* BL21-DE3-RP cells. Corresponding nPIST fragments were purified and run in SDS PAGE (Fig. 5B, 5C, 5D, and 5E). Like full length nPIST and ΔWH2 fragment (268^th^ - 463^rd^ aa), the PDZ fragment (285^th^ – 463^rd^ aa) also occurred in higher molecular size (~27 KDa) than its theoretical value 22.7 KDa with 6-His tag in SDS-PAGE (Fig 5E). The WH2 fragment (200^th^ – 289^th^ aa) protein was small in size and very degradation prone. So, it could not be further purified by gel-filtration chromatography to get rid of non-specific bands (Fig. 5D). The CC1, CC2, WH2 and PDZ fragments were subjected to F-actin co-sedimentation assay, the supernatant and pellet fractions separated and run in SDS PAGE. The co-sedimentation assay gel image of CC1 fragment showed that the band intensity of CC1 in pellet fraction was present in rising order whereas the pellet fraction of only 12 μM CC1 negative control reaction had little CC1 fragment protein (Fig. 5F). The co-sedimentation assay of CC2 fragment (Fig. 5G) and WH2 fragment (Fig. 5H) exhibited their presence in pellet fraction in escalating amount when incubated with 5 μM F-actin in concentration dependent manner. On contrary, the pellet fractions of the only CC2 fragment and WH2 fragment reaction had very negligible concentration of corresponding constructs (Fig. 5G and H). We also found that PDZ fragment (285^th^ – 463^rd^ aa) got co-sedimented with actin in pellet fraction in negligible amount (Fig. 5I). Here to be noted that one actin binding region is lying between WH2 domain and PDZ domain (268^th^ – 284^th^ aa) which can be indicated from rapid reduction in affinity for actin of PDZ fragment (Fig. 5I) upon deletion of these amino acids from ΔWH2 fragment ( Fig. 4D) but identification amino acids responsible for actin interaction needs further investigation.

**Figure 5:**
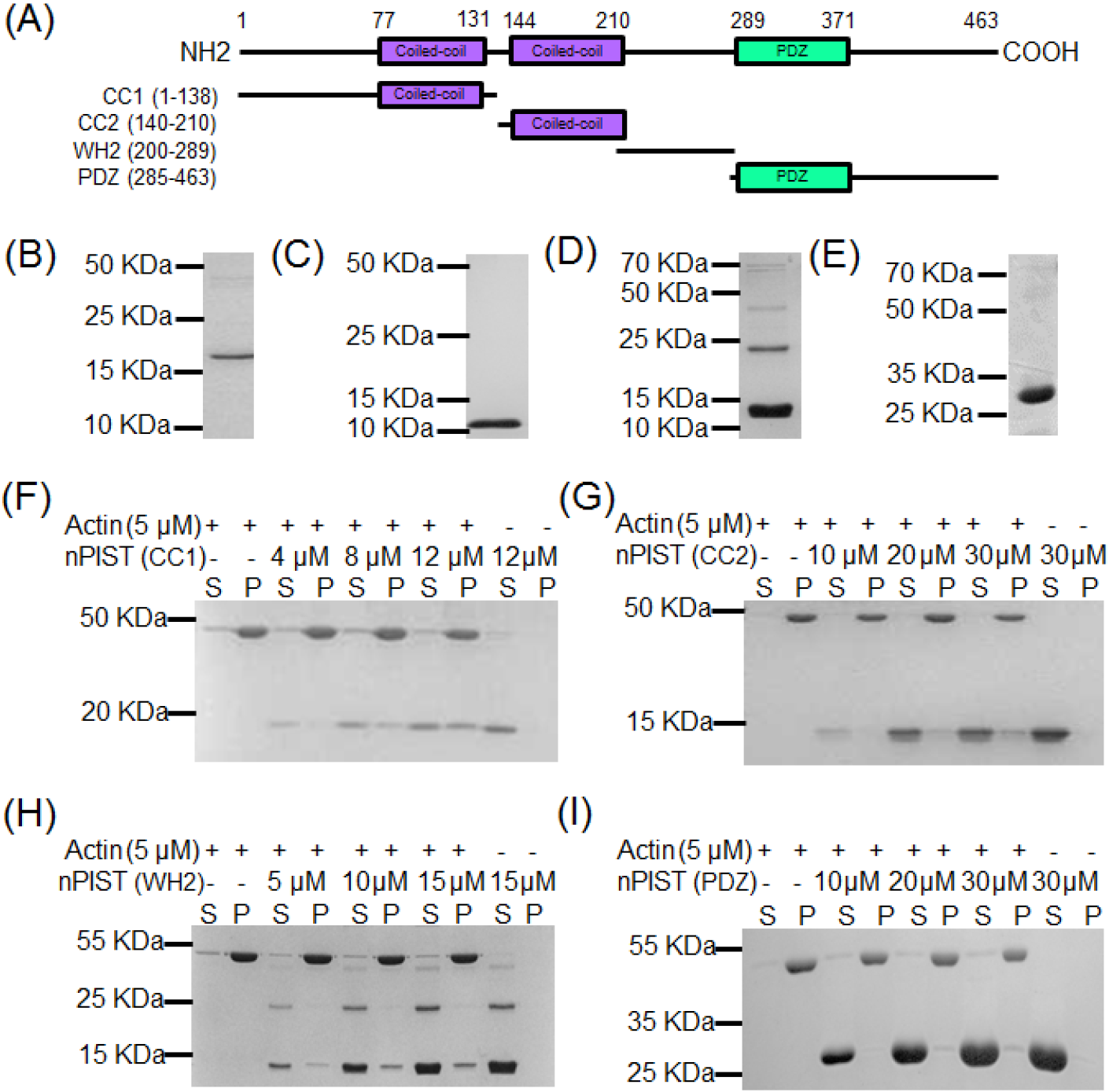
Presence of several actin interacting regions makes nPIST an actin binding protein. (A) Schematic diagram of truncated fragments of nPIST considered to identify actin binding regions. 6-His tagged purified protein of (B) CC1 fragment (1^st^ – 138^th^ aa), (C) CC2 fragment (140^th^ – 210^th^ aa), (D) WH2 fragment (200^th^ −289^th^ aa), and (E) PDZ fragment (285^th^ −463^rd^ aa) of nPIST run in coomassie stained 12% SDS-PAGE. Gel electrophoresis of supernatant and pellet fraction of F-actin co-sedimentation assay reactions of (F) CC1 fragment, (G) CC2 fragment, (H) WH2 fragment, and (I) PDZ fragment of nPIST.

### Full length nPIST unable to bundle actin filaments

npist having multiple actin binding motifs, we hypothesized that it might act as a filamentous actin bundling protein. To study that, we incubated 1 μM, 2 μM and 3 μM of nPIST with 5 μM actin and the reactions are centrifuged at much lower speed; 10,000 rpm (Shimada *et al.*; 2004). Human Fascin1 was used as a positive control. The supernatant and pellet fractions of each reaction were run in SDS PAGE. It displayed that most of the actin when incubated with nPIST was present in supernatant fraction like that of actin control reaction whereas actin majorly appeared in the pellet fraction in presence of positive control Fascin (Maekawa *et al.*; 1982). nPIST was also mainly present in supernatant fraction along with the actin (Fig. S1A). It indicated that full length nPIST did not bundle actin filaments.

### nPIST acts as actin stabilizing agent

We further analyzed the effect of full length nPIST on *in vitro* actin dynamics. In dilution dependent depolymerization assay, 50% pyrene-labeled preassembled F-actin filaments diluted to 0.1 μM concentration were used. The pyrene fluorescence gradually decreased with time which denoted depolymerization of F-actin filaments (Fig 6A and S1B). In presence of nPIST, prevention of depolymerization of F-actin filament is indicated by reduction in fluorescence decrease rate (Fig 6A and B). *Xenopus* cofilin1 (Xac1) was used as negative control (Rosenblatt *et al.*; 1997) and when incubated with F-actin, it severed the F-actin filaments which can be visualized by rapid drop in fluorescence intensity. When nPIST was provided in the reaction, it inhibited the F-actin (diluted to 0.25 μM) severing effect of Xac1 in a concentration dependent manner (Fig S1B). We also tested the effect of nPIST on depolymerisation of F-actin filaments diluted to 0.25 μM but there was not much change from actin control reaction (Fig. S1C). This notified that full length nPIST stabilized F-actin filaments *in vitro* and inhibited Xac1 mediated F-actin severing.

**Figure 6:**
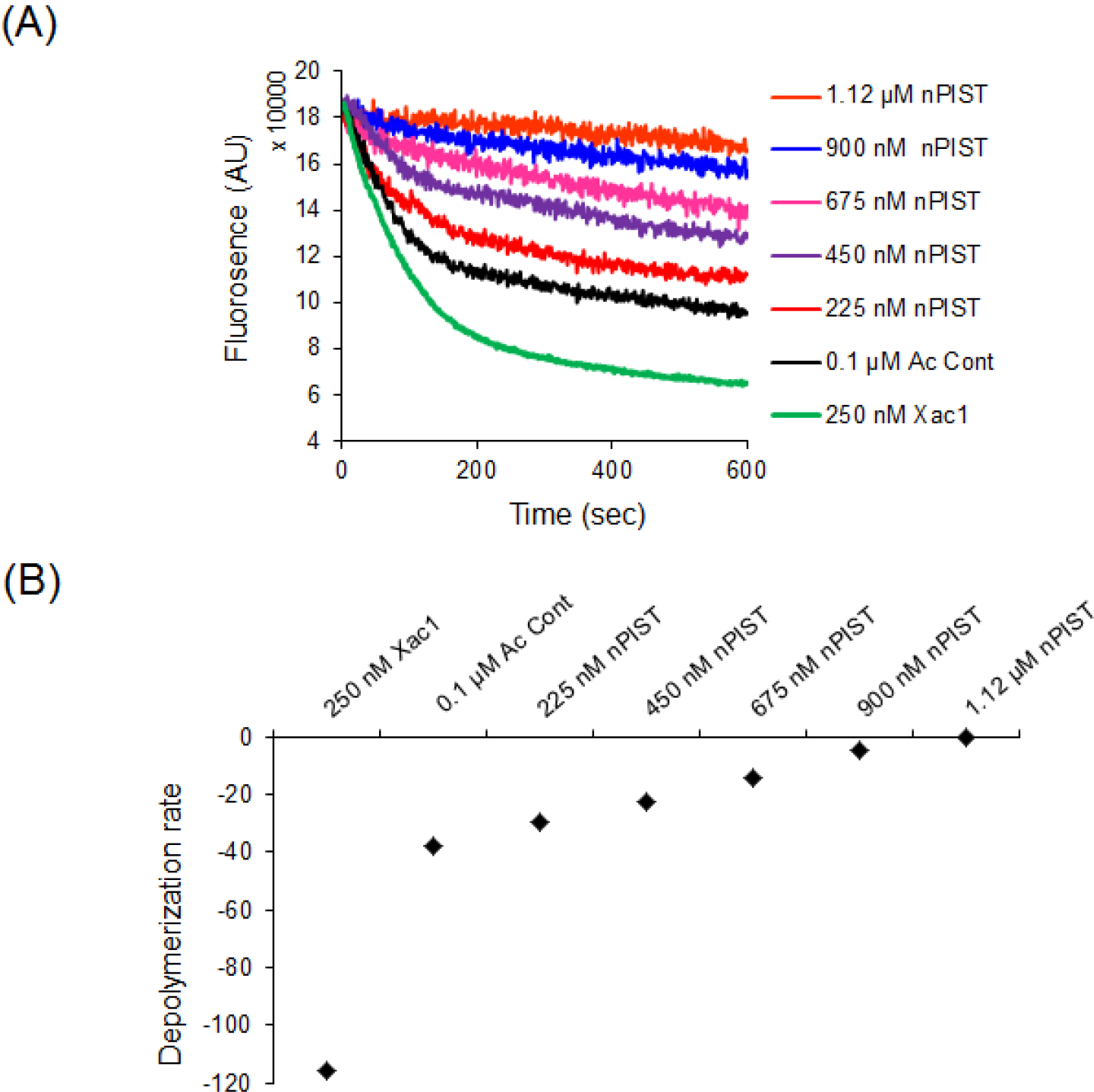
nPIST stabilizes F-actin filaments *in vitro*. (A) F-actin filament stabilization assay with 0.1 μM 50% pyrene-labeled actin in presence of increasing concentration of nPIST. *Xenopus* cofilin 1 (Xac1) used as filament stabilization negative control and severed actin filaments marked by the rapid decrease in fluorescence intensity of N-pyrene. (B) Actin filament stabilization rate measured as the slope of fluorescence curve for 10-100 secs.

### nPIST causes unusual accumulation of cellular actin through CC-WH2 domain

Next, we looked into the effect of nPIST and its fragments on *in vivo* actin cytoskeleton. N-terminal GFP-tagged full length nPIST, CC-WH2 (1^st^ – 267^th^ aa), and ΔWH2 (268^th^ – 463^rd^ aa) fragment were transiently transfected in cultured HEK293 cells (Fig. 7). Cells were fixed 36 hours post transfection and actin was visualized by rhodamine phalloidin staining (1^st^ column). GFP and rhodamine channel merged (3^rd^ column) images showed that nPIST was primarily localized in the whole cytoplasm as well as got confined near nucleus to a great extent (3^rd^ row). The expression of CC-WH2 fragment (4^th^ row) was almost exclusively restricted to perinuclear region of the cell but ΔWH2 fragment which did not have golgi localization region (5^th^ row) lost perinuclear expression pattern. Interestingly, nPIST and CC-WH2 fragment co-localized with a patch of actin. On the other hand ΔWH2 fragment did not show such effect on actin cytoskeleton which was similar to that of only GFP expression (2^nd^ row). From the above experiment we could suggest that transient over expression of nPIST causes abnormal clustering of F-actin inside cultured HEK293 cells.

**Figure 7:**
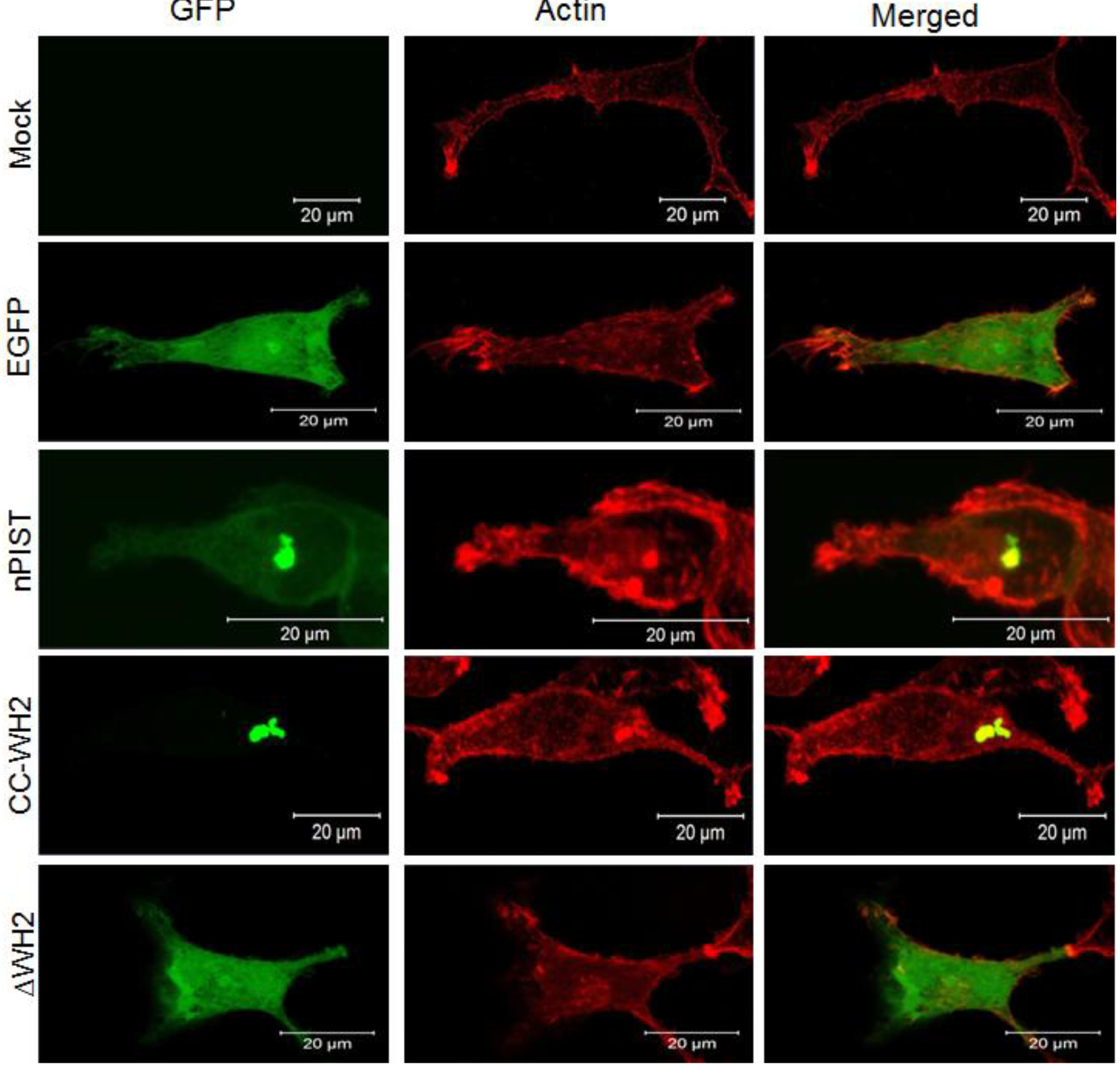
Effect of Full Length nPIST and its Fragments on Cellular Actin Cytoskeleton. Full length nPIST and its fragments (CC-WH2 and ΔWH2) was sub cloned in pEGFP-C1 vector. HEK 293 cells were transfected with GFP vector control (2^nd^ row), full length nPIST (3^rd^ row), CC-WH2 fragment (4^th^ row), and ΔWH2 fragment (5^th^ row). Actin was stained with rhodamine-phalloidin (1^st^ column). Third column shows merged actin and GFP.

## Discussion

In the current study, we have depicted nPIST as a novel actin binding protein. In silico model of nPIST WH2 domain (233^th^ – 249^th^ aa) and G-actin complex implemented that nPIST may bind to actin adequately (Fig. 1E). Our experimental data validated the in silico data and for the first time we report nPIST acting as an actin binding protein (Fig 2C). Further sequence based analysis of the protein disclosed that nPIST contains several other motifs which act as actin binding site for the protein. We indicated the presence of another three actin binding regions in coiled-coil helix 1 region (CC1, 1^st^ – 138^th^ aa), coiled-coil helix 2 region (140^th^ – 210^th^ aa) and C-terminal PDZ domain region (ΔWH2, 268^th^ – 463^rd^ aa) (Fig. 4A, and 5A). F-actin co-sedimentation assay results showed that each of these fragmented regions can individually bind to actin (Fig. 4D, 5F, 5G, and 5H). However, to determine the exact amino acids residues involved in actin binding needs further detailed study. The presence of several actin binding regions makes nPIST capable of binding to actin. Our experimental data also showed that nPIST can stabilize F-actin filaments *in vitro* and inhibited filament severing effect of Xac1 (Fig. 6A, 6B, S1B and S1C). We have also investigated the effect of nPIST on actin cytoskeleton *in vivo* (Fig. 7). Full length nPIST and ΔWH2 (268^th^ - 463^rd^ aa) was found to be expressing throughout the cell cytosol (Fig. 7, 3^rd^ and 5^th^ row) whereas CC-WH2 (1^st^ – 267^th^ aa) was exclusively localized to a perinuclear region of cytoplasm (Fig. 7, 4^th^ row). nPIST and CC-WH2 fragment co-localized with phalloidin stained actin patch but such phenomenon was not observed in case of ΔWH2 fragment. Occurance of similar kind of F-actin accumulation can also be seen in case of overexpression of Slingshot phosphatase (Soosairajah *et al.*, 2005) but not in juxtanuclear region. The mechanism behind the occurrence of such actin patch needs further inspection to unfold the underlying story. Strikingly, nPIST and the CC-WH2 fragment consisting of golgi localization region in the second coiled-coil helix region got spatially localized to the juxtanuclear region. On the other hand, the ΔWH2 fragment lacking golgi localizing region lost its targeted expression and got distributed in the whole cytosol. Whether the co-localization of nPIST and actin was in golgi region or not can be disclosed by co-localization study with golgi resident markers. The actin binding regions were mostly present within CC-WH2 fragment and nPIST was capable of co-localizing actin through its CC-WH2 fragment but ΔWH2 fragment having only one of such stretch interacting with actin domain did not exert much any effect on cellular actin cytoskeleton. Till date studies have suggested that WASP family proteins have evolved to mostly promote Arp2/3 complex mediated actin nucleation (Kollmar *et al.*; 2012). But some members of this family have Arp2/3 independent activity and directly function as actin nucleating candidate (Dominguez *et al.*; 2016). Our candidate protein nPIST is another WH2-like domain containing protein that directly interacts with actin and modulates its dynamics *in vitro*.

The proper positioning and maintenance of golgi morphology requires presence of actin microfilaments (Valderrama *et al.*; 1998). Several actin binding proteins located in golgi network have been reported to be linking actin cytoskeleton to vesicular trafficking regulation, modeling of golgi architecture and integrating molecular machinery for vesicular transport (Egea *et al.*; 2006). Among these microfilament regulators, there are several Wiskott–Aldrich syndrome protein (WASP) family candidates involved in different compartments of vesicle mediated transport pathway. WHAMM is known to regulate trafficking in ER-Golgi intermediate compartment (ERGIC) compartments (Kenneth et al.; 2008). The closest family member of WHAMM; JMY on the other hand is mainly involved in the trans-Golgi network (TGN) anterograde trafficking (Schlüter et al.; 2014). Another WASP family member WASH functions in regulation of endosome shape and fission (Derivery et al.; 2009, Duleh and Welch; 2010). The molecular mechanism underlying nPIST mediated vesicular trafficking regulation in terms of actin cytoskeleton regulation remains to be elucidated which can be an important functional aspect to be explored that can have considerable contribution in the broad spectrum of purpose served by WASP family proteins (Alekhina *et al.*; 2017).

## Materials and methods

### Computational Analysis of nPIST WH2 domain

The structural model of nPIST was obtained using Swiss model and modeller (Kelley *et al.*; 2009). Sequences of WH2 domain from all the crystal structure available were extracted and aligned with the sequence of nPIST, resulted in identification of WH2 domain in the region (233^rd^ -249^th^ aa) of nPIST. The model of G-actin-WH2 domain complex was obtained using the cross-linked complex of actin with first WH2 domain of *V. parahaemolyticus* VopL (PDB ID: 3M1F) (Reboeski *et al.*; 2010). The WH2-domain was docked using rosetta Flexpepdock server (London *et al.*; 2011). The starting structure was refined in 200 independent FlexPepDock simulations. 100 of simulations were carried out strictly in high-resolution mode, while 100 of simulation included a low-resolution pre-optimization step, followed by high-resolution refinement. A total of 200 models were created and then ranked based on their Rosetta generic full-atom energy score. The final model of the complex was further energy minimized using AMBER12 molecular dynamics software (Case *et al.*; 2012). Analysis of the complex using KFC-2 server (Zhu *et al.*; 2011), predicted binding “hot spots” within protein-protein interfaces by recognizing structural features indicative of important binding contacts. The figures were prepared using Pymol software (The PyMOL Molecular Graphics System, Version 1.5.0.4 Schrödinger, LLC).

### Plasmid constructs

Total RNA was isolated using TRIzol (Invitrogen) from adult brain tissue of C57BL/6 mouse; cDNA was prepared by RT-PCR (Superscript III, Invitrogen), and full length sequence was amplified with forward 5’GGAAGATCTATGTCGGCGGGTGGC3’ and reverse 5’CCGCGTCGACTTAGTAGGCCTTCTTCTGATGCA3’ primers. Full length nPIST was cloned in pET-28a vector (Novagen) using BamHI and SalI, expressed in *E. coli* BL21-DE3-RP as N-terminal 6-His tagged protein. The sub clones of nPIST fragments were prepared in similar manner. The primer used for sub-cloning were; forward 5’GGAAGATCTATGTCGGCGGGTGGC3’ and reverse 5’ CCGCGTCGACTTACATGGGCCGCTTCAAG 3’ for CC-WH2 (1^st^ – 267^th^ aa) fragment, forward 5′GGAAGATCTCAAGCACCCCCAGGCC3′ and reverse 5’CCGCGTCGACTTAGTAGGCCTTCTTCTGATGCA3’for ΔWH2 (268^th^ – 463^rd^ aa) fragment, forward 5’GGAAGATCTATGTCGGCGGGTGGC3’ and reverse 5′CCGCGTCGACTTAAGTCTTGGCATGAAGCTGAAG3′ for CC1 (1^st^ – 138^th^ aa) fragment, forward 5′GGAAGATCTCAAAGTGTTGACTCTGGGG3′ and reverse 5′CCGCGTCGACTTAGTACTTGGCAGCCAACCTC3′ for CC2 (140^th^ -210^th^ aa) fragment, forward 5′GGAAGATCTGAAGTTTATGGGGCGAGG3′ and reverse 5’ CCGCGTCGACTTATTTTCTAATTGGACCGACTC3’ for WH2 (200^th^ – 289^th^ aa) fragment, and forward 5’ GGAAGATCTCCAATTAGAAAAGTTCTCCTCC 3’ and reverse 5’ CCGCGTCGACTTAGTAGGCCTTCTTCTGATGCA3’ for PDZ (285^th^ – 463^rd^ aa) fragment. Human Fascin1 construct was a kind gift from Professor Jo Adams and Dr. Aurnab Ghose. It was sub-cloned in pET28a vector using forward 5’ GACGAGGATCCATGACCGCCAACGGCACAG3’ and reverse 5’ CAGCTCGAGCTAGTACTCCCAGAGCGAGGCG3’ primers.

### Protein Purification

G-actin was purified from rabbit muscle acetone powder (Pollard TD; 1984) with G-Buffer [5 mM Tris pH 8.0 (Sigma-Aldrich), 0.2 mM ATP (USB), 0.2 mM CaCl_2_ (USB) and 0.2 mM DTT (USB)]. Actin was taaged with N-(-1-pyrene) iodoacetamide (P-29, Molecular probe) for fluorescence spectroscopic analysis (Higgs and Pollard, 1999). *E. coli* BL21-DE3-RP (Stratagene) containing nPIST, its different fragment constructs and Fascin1 were grown up to O.D 0.6 at 37°C, induced with 0.5 mM IPTG (Thermo Fisher Scientific), harvested, and stored in −80°C (Moseley *et al.*; 2006). Harvested cells were resuspended in lysis buffer [50 mM Tris-Cl pH 8.0, 100 mM NaCl (Sigma-Aldrich), 30 mM Imidazole pH 8.0 (USB), 0.5 mM DTT, 0.5% IGEPAL (Sigma-Aldrich), and Protease Inhibitor Cocktail], lysed by sonication for 5 minutes in 1 minute pulse in 30 second interval, and then centrifuged at 12000 rpm. The supernatant was incubated with 50% slurry of Ni-NTA resin beads (Qiagen), washed with wash buffer (50 mM Tris-Cl pH8.0, 150 mM NaCl, 50 mM Imidazole pH8.0) and finally eluted with elution buffer [50 mM Tris-Cl pH8.0, 100 mM NaCl, 350 mM Imidazole pH 8.0, and 5% glycerol (Invitrogen)]. Next, purified cloned constructs were dialyzed in HEKG_5_ buffer [20 mM HEPES (USB), 1 mM EDTA (Calbiochem), 50 mM KCl (Sigma-Aldrich), and 5% Glycerol) (Moseley *et al.*; 2006). nPIST and its fragments were further purified using Superdex-200 10/300 GL column (GE Healthcare).

*Xenopus* cofilin 1 in pGEX_(GST-Xac1) was kindly gifted by Prefessor JR Bamburg (Abe *et al.*; 1996). GST-tagged Xac1 was expressed in E. coli BL21-DE3 (Stratagene) at 37oC and induced with 0.5 mM IPTG for 4 hours. Post harvestion, cells were resuspended in lysis buffer (50 mM Tris-Cl pH 8.0, 150 mM NaCl, 1 mM DTT, 1 mM DTT, 0.5% IGEPAL, and Protease Inhibitor Cocktail), sonicated, incubated with 50% slurry of GST-Agarose bead (thermo Scientific) and eluted with elution buffer having 50 mM Tris-Cl pH 8.0, 150 mM NaCl, 1 mM EDTA, and 10 mM Reduced Glutathione (USB).

### Co-sedimentation Assay

5μM G-actin was polymerized into F-actin for 1 hour at 25°C in F-buffer [10 mM Tris-Cl pH 8.0, 0.2 mM DTT, 0.5 mM ATP, 50 mM KCl, and 2 mM MgCl_2_ (Sigma-Aldrich), 0.2 mM CaCl_2_]; subsequently nPIST was added for 15 minutes and then centrifuged at 310 x 1000 g for 40 minutes in TLA-100 rotor (Beckman Coulter). Supernatant was separated from pellet fraction and pellet was resuspended in F-buffer. All the samples were boiled with 4X sample loading buffer, loaded on SDS-PAGE, visualized with coomassie (SRL) staining. The above procedure was carried out for different fragments of nPIST to check their actin binding property (Shimada *et al.*; 2004).

### Interaction kinetics by Surface Plasmon Resonance (SPR)

SPR experiments were performed using a Biacore T200 system. All experiments were conducted in duplicates at 25°C and double referencing (blank immobilized channel and blank injections) was used to eliminate bulk effects. nPIST (ligand) was immobilized on a series S CM5 sensor chip using amine coupling to achieve an immobilization level of ~1500 RUs. The running buffer was varied according to the analyte being used (G-buffer for G-actin and F-buffer for F-actin). Varying concentrations of the analytes (0.5 μM, 1 μM, 2 μM, 4 μM, and 8 μM) were passed over the immobilized ligand at a flow rate of 10 μl/min for 180 s to observe the association phase and the dissociation was observed by flowing blank buffer for 120 s. Data analyses were carried out with Biacore T200 evaluation software 2.0 to fit the data to 1:1 interaction model.

### F-actin depolymerization assay

5 μM of 50% pyrene-labeled rabbit muscle actin was incubated in F-buffer and polymerized into F-actin for one hour at 25° C. From this preassembled actin, 2 μl was used in 100 μl of reaction having 83 μl of F-buffer and 15 μl of HEKG5 buffer or protein samples. The N-pyrene (Invitrogen) fluorescence was excited at 365 nm and the emitted spectrum was measured at 407 nm for 600 seconds (QM 40 PTI NJ) (Rosenblatt *et al.*; 1997).

### Cellular Experimentation

HEK 293 cells (ATCC®-CRL-1573) were grown in poly D-lysine coated cover slips in 24-well plate with MEM (Gibco) containing L-Glutamine (Gibco), penicillin streptomycin (Hyclone, Thermo Scientific) and 10% Fetal Bovine Serum (Gibco). Full length nPIST and its CC-WH2 (1^st^ – 267^th^ aa) and ΔWH2 (268^th^ – 463^rd^ aa) fragments were cloned in pEGFPC1 vector (Stratagene). Cells were transfected with cloned nPIST constructs with Lipofectamine 2000 (Invitrogen) following manufacturer’s protocol. The cells were fixed using 4% paraformaldehyde (Sigma-Aldrich), 36 hours post transfection. Cellular actin was visualized with 50 nM rhodamine phalloidin staining (Invitrogen) and nPIST and its fragments were ectopically expressed as N-terminal GFP tagged proteins. The cell containing cover slips were mounted with antifade reagent (Thermo Fisher Scientific) and images were captured with laser scanning microscope (Carl Zeiss LSM710) using 488 laser and 561 laser.

## Acknowledgements

SD thanks IISER-Kolkata and MM thanks ICMR for providing their fellowship. Xac1-GST was gifted to SM by Prof JR Bamburg. Human Fascin1 cloned in pEGFP-C1 was a kind gift from Professor Jo Adams and Dr. Aurnab Ghose. This work is supported by IISER Kolkata.

## Authors’ contrubutions

Designed and conceived the experiment: SM, PD, SD, SS, KD, DS, MM and SG. Experiments performed by: PD, SD, MM, SS, KD. Analyzed the data: SM, PD, SD, SS, KD, DS, MM and SG. Writing of the manuscript: SM, PD, SD, SS, KD, DS, MM and SG.

---

**Figure S1:**
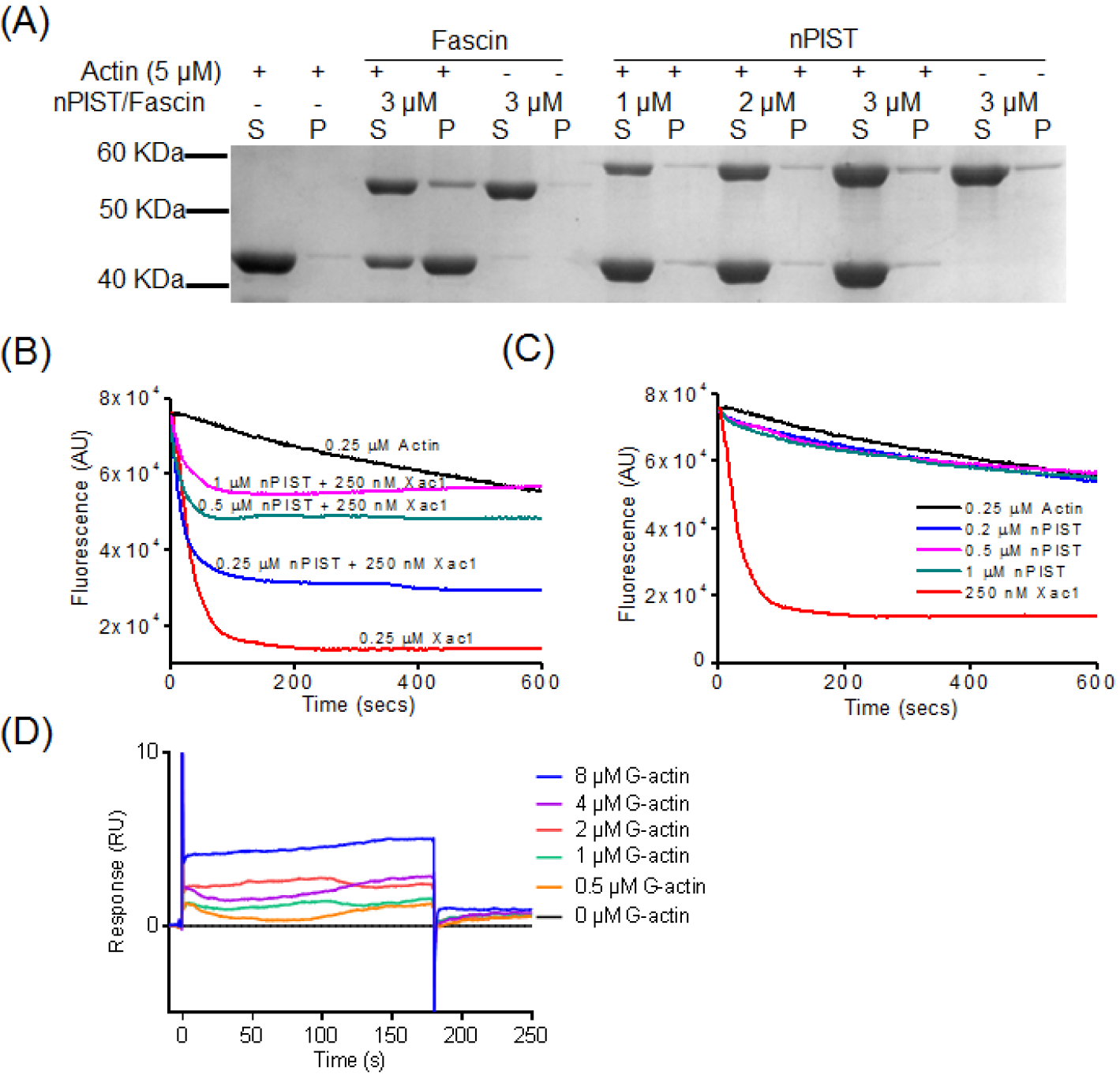
Full length nPIST does not bundle but stabilizes actin filaments. (A) *In vitro* low speed F-actin co-sedimentation assay of full length nPIST performed to check its bundling ability. 5 μM actin and 1, 2 and 3 μM of nPIST incubation reaction was centrifuged at 9.2 x 1000 g for 10 minutes at 4°C and supernatant and pellet fraction run in SDS-PAGE is denoted by S and P respectively. 3 μM human Fascin1 was used as positive control. (C) Actin filament stabilization assay with 0.25 μM of 50% pyrene-labeled actin. nPIST salvaged the actin filaments from being severed by negative control Xac1 in dose dependent manner. (D) Dilution dependent depolymerization assay of only 0.25, 0.5 and 1 μM of nPIST with 0.25 μM of actin where Xac1 was used as negative control. (E) SPR sensorgram for the binding of G-actin to immobilized nPIST, showing the instable interaction.

